# Hierarchical representation of multi-step tasks in multiple-demand and default mode networks

**DOI:** 10.1101/582858

**Authors:** Tanya Wen, John Duncan, Daniel J Mitchell

## Abstract

Task episodes consist of sequences of steps that are performed to achieve a goal. We used fMRI to examine neural representation of task identity, component items, and sequential position, focusing on two major cortical systems – the multiple-demand (MD) and default mode networks (DMN). Human participants (20 male, 22 female) learned six tasks each consisting of four steps. Inside the scanner, participants were cued which task to perform and then sequentially identified the target item of each step in the correct order. Univariate time-course analyses indicated that intra-episode progress was tracked by a tonically increasing global response, plus an increasing phasic step response specific to MD regions. Inter-episode boundaries evoked a widespread response at episode onset, plus a marked offset response specific to DMN regions. Representational similarity analysis was used to examine encoding of task identity and component steps. Both networks represented the content and position of individual steps, but the DMN preferentially represented task identity while the MD network preferentially represented step-level information. Thus, although both DMN and MD networks are sensitive to step-level and episode-level information in the context of hierarchical task performance, they exhibit dissociable profiles in terms of both temporal dynamics and representational content. The results suggest collaboration of multiple brain regions in control of multi-step behavior, with MD regions particularly involved in processing the detail of individual steps, and DMN adding representation of broad task context.

**Significance Statement:** Achieving one’s goals requires knowing what to do and when. Tasks are typically hierarchical, with smaller steps nested within overarching goals. For effective, flexible behavior, the brain must represent both levels. We contrast response time-courses and information content of two major cortical systems – the multiple-demand (MD) and default mode networks (DMN) – during multi-step task episodes. Both networks are sensitive to step-level and episode-level information, but with dissociable profiles. Intra-episode progress is tracked by tonically increasing global responses, plus MD-specific increasing phasic step responses. Inter-episode boundaries evoke widespread responses at episode onset, plus DMN-specific offset responses. Both networks encode content and position of individual steps, but the DMN and MD networks favor task identity and step-level information respectively.

## Introduction

Purposeful behavior requires retrieval of memorized sequences (Hsieh and Ranganath, 2015) to guide current actions, with overarching goals or “task episodes” (e.g., “make stew”) decomposed into achievable steps (“wash vegetables” → “chop” → “cook”) (Cooper and Shallice, 2000; Schneider and Logan, 2006; Duncan, 2010; Farooqui and Manly, 2019). As each step is completed, its specific content loses relevance, while higher-level representations of the full task episode remain in behavioral control. This raises the question of how brain regions work together to execute a current step while keeping an overall goal in mind.

Previous literature highlights the importance of a frontoparietal “multiple-demand” (MD) network in controlling complex mental programs (Duncan, 2010, 2013). MD regions are sensitive to hierarchical task structure (Farooqui et al., 2012; Desrochers et al., 2015), and are well-suited for focusing on specific contents of current cognition, preferentially encoding task-relevant information (Asaad et al., 2000; Everling et al., 2002; Li et al., 2007), and radically changing activity patterns across successive task steps (Sigala et al., 2008).

Representation of event sequences is also critical in episodic memory research (Ezzyat and Davachi, 2011; Hsieh et al., 2014; Cohn-Sheehy and Ranganath, 2017). Event segmentation theory (Zacks and Tversky, 2001; Radvansky and Zacks, 2017) proposes that humans segment incoming information into temporally meaningful chunks; when important situation features change, an update to the event model is experienced as an event boundary. Event boundaries may activate regions resembling the MD network (Zacks et al., 2001; Sridharan et al., 2007), but also areas associated with episodic memory, including the hippocampus (Ben-Yakov et al., 2013; Ben-Yakov and Henson, 2018) and the default mode network (DMN) (Speer et al., 2007), or both (Ezzyat and Davachi, 2011). The DMN is implicated in high-level cognition at a broad scale, including encoding of schemas (Robin and Moscovitch, 2017), situation models (Reagh and Ranganath, 2018), and cognitive contexts (Crittenden et al., 2015). The DMN response to event boundaries likely reflects meaningful extended episodes, because activation is greater at boundaries rated as separating long meaningful events than boundaries separating shorter events (Speer et al., 2007). Similarly, temporal scrambling of narrative stimuli suggests a cortical hierarchy of temporal receptive windows (Lerner et al., 2011), with short-timescale processing in sensory regions, where inter-subject pattern consistency is robust to fine-scale scrambling, intermediate timescales in MD regions, and longest-timescale processing in DMN regions, where pattern consistency is sensitive to even coarse scrambling (Chen et al., 2016). An investigation of multi-voxel pattern transitions during narrative perception (Baldassano et al., 2018) found the longest-timescale event representations in posterior medial cortex and the intraparietal sulcus, within the DMN and MD networks respectively. Neural event structure was found to match across visual and auditory modalities near the temporoparietal junction and in lateral frontal cortex, again within DMN and MD networks respectively. Overall, the literature suggests that both DMN and MD networks are potentially well-suited to representing task episodes over extended timescales.

To our knowledge, no study has contrasted the roles of the DMN and MD networks in representing different aspects of task episodes. We therefore examined how these networks represent information at multiple levels of abstraction within a task: individual steps, including content and position within an episode, the whole task, and groups of related tasks. Participants learned four-step tasks associated with different rooms (kitchen or bathroom), and then sequentially identified target items corresponding to each step of cued tasks. Thus, we could quantify neural representation of rooms (e.g. kitchen), tasks (e.g. “make stew”), steps (e.g. third) and items associated with steps (e.g. “wash vegetables”). We used univariate event-related and finite impulse response (FIR) analyses to characterize the temporal evolution of activity across episodes, and representational similarity analysis (RSA) to investigate representations of task structure and content. We hypothesized that different brain regions would be preferentially sensitive to different levels of the temporal task hierarchy, and focused on the MD and DMN networks as *a priori* regions of interest.

## Methods

### Participants

42 participants (20 male, 22 female; ages 18-39, mean = 26.79, SD = 4.77) were included in the experiment at the MRC Cognition and Brain Sciences Unit. An additional 19 participants were excluded (two were discovered to have cysts, one lost several slices due to poor bounding box positioning, ten were excluded due to having no correct episodes for at least one combination of cued task × distractor task (see later), and a further six were excluded due to excessive head motion > 5 mm). All participants were neurologically healthy, right-handed, with normal or corrected-to-normal vision. Procedures were carried out in accordance with ethical approval obtained from the Cambridge Psychology Research Ethics Committee, and participants provided written, informed consent before the start of the experiment

### Stimuli and task procedures

The study consisted of a learning session outside the scanner and an execution session in the scanner. During the learning session, participants learned six everyday task sequences, each based in one of two locations (“rooms”; three kitchen and three bathroom). Each task consisted of four ordered “steps”. For example, the task “make a stew” consisted of the steps “take food from fridge”, “wash vegetables”, “chop vegetables”, “cook on stove”. Each step was associated with a unique image (“item”). The complete set of stimuli is shown in Figure 1A.

**Figure 1.**
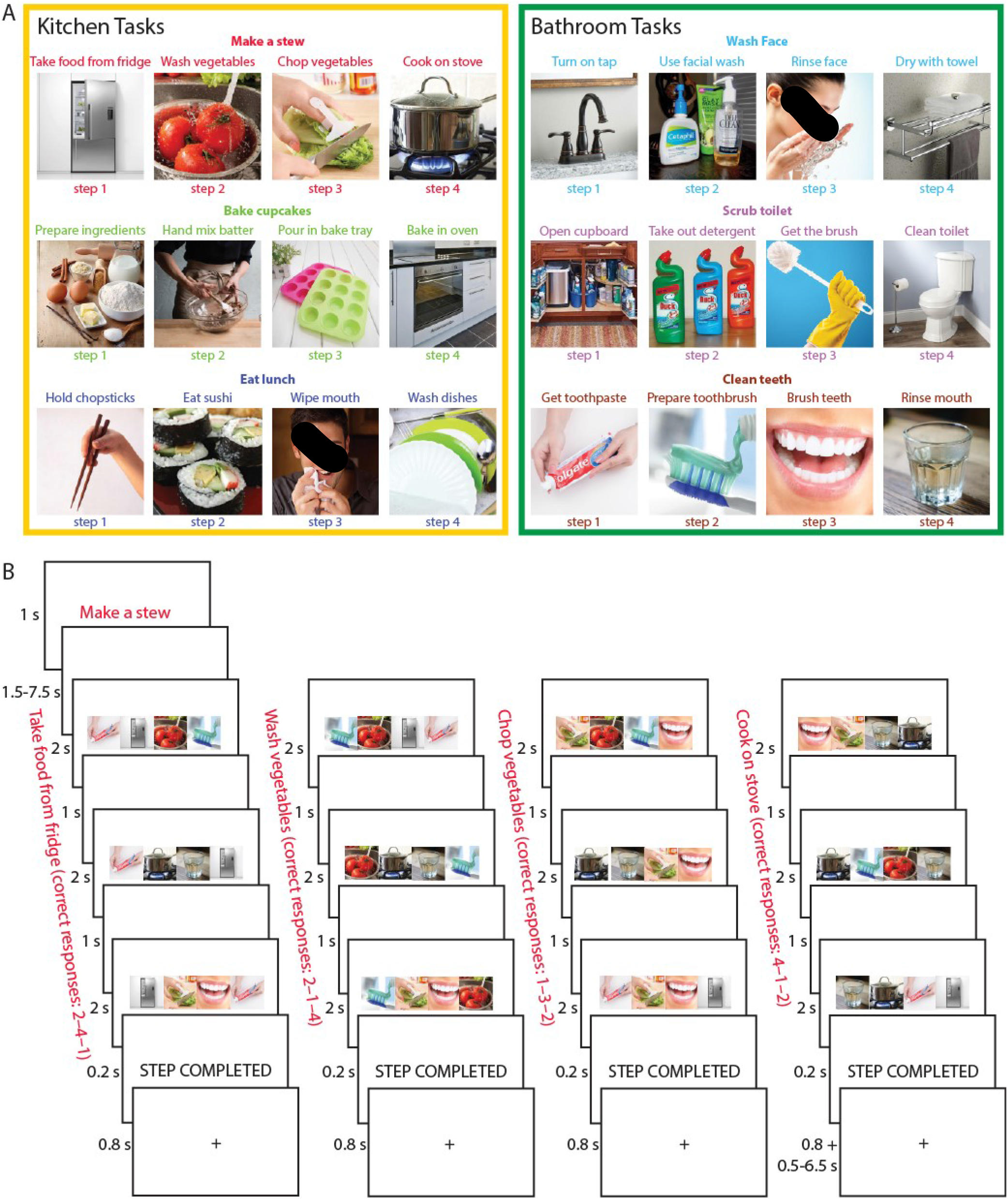
(A) Illustration of the six tasks (three kitchen and three bathroom tasks) memorized before going into the scanner. Each task consisted of four steps to be completed in serial order (e.g., the task “make a stew” consisted of “take food from fridge”, “wash vegetables”, “chop vegetables”, “cook on stove”). (B) Structure of an example task episode. Episodes began with a cue indicating which task to perform (e.g., “make a stew”). After a short delay, the first search array of four items appeared, and participants were asked to select the item corresponding to the first step of that task (here, “take food from fridge”). Participants selected this same target in three search arrays (total step duration = 9 s), then were given a brief indicator that the step had been completed, and moved on to the next step (here “wash vegetables”). Completion of all four steps completed the entire task episode.

In the learning session, participants viewed the names and images of the steps of each task in sequential order. The step images were presented simultaneously with a background image corresponding to the room in which they occur (kitchen or bathroom). The learning was self-paced, in separate runs for each room. Within each room, each task sequence was presented three times, and each item within the sequence was presented until the participant decided to move on to the next item. There was a 1.5 s inter-stimulus interval between items. After viewing all six sequences, participants were tested for their memory of the tasks by (1) sorting picture cards representing all steps of the six tasks into the correct sequences, and (2) completing a pen-and-paper test in which they were asked to write down the names of the steps in the correct order for each task. Most participants performed both tests without error. A few participants made a mistake on 1-2 items but were able to correct their answers after being told they made a mistake. The tests ensured participants had memorized the specific step sequence of each task. Before entering the scanner, participants practiced a shortened version of the main experiment, containing one episode of each task. During scanning, participants performed two runs of the experiment, interleaved with shorter runs (~5 minutes) of a localizer task that was not analyzed and is not described further.

Figure 1B illustrates the structure of the task episodes paradigm. At the start of each 45 s episode, participants were presented with a cue (e.g., “make a stew”) for 1 s, indicating which task to complete. This was followed by a fixation period lasting between 1.5 – 7.5 s, selected randomly from a uniform distribution, before the onset of the first step. On each step, participants had to perform three visual searches. On each search, an array of four images was presented in a horizontal row (total left to right visual angle approximately 12.6°). These included (randomly ordered from left to right): (1) the correct image (“target”) corresponding to the current task step; (2) a distractor image representing a random incorrect step from the correct task; (3) a distractor representing the correct step but from an incorrect task (“distractor task”); and (4) an additional distractor representing the same incorrect step as (2), from the same incorrect task as (3). To ensure that each display contained two images from each room, distractor tasks were selected at random from the alternative room to the cued task. The array remained for 2 s, and within this time, the participant had to indicate the position of the target image using a 4-choice button box with their right hand. A 1 s fixation interval preceded onset of the next search array. Each step thus lasted for 9 s, with the participant selecting the same target in each of three search events, to allow separation of the hemodynamic response to successive task steps, while ensuring sustained focus on the relevant item within each step. At the end of the third search event, a 0.2 s presentation of the words “STEP COMPLETED” indicated the completion of that step, followed by a 0.8 s fixation interval. Without further cueing, the participant then moved on to the next task step. After completing the last step, a fixation interval of 0.5 – 6.5 s was presented before the onset of the cue for the next task. The total interval between the last step of the previous task and the first step of the next task was fixed at 9 s. Participants were not given feedback on their accuracy. Each run consisted of 36 task episodes (with an additional dummy episode to start), constructed so that each task appeared following each possible preceding task once. Task ordering was chosen before the start of each run to maximize the design efficiency (Dale, 1999) of all pairwise contrasts between tasks. 1000 task orders were simulated, and the most efficient one was chosen. Each of the two runs lasted ~28 min.

### fMRI data acquisition and preprocessing

Scanning took place in a 3T Siemens Prisma scanner. Functional images were acquired using a multi-band gradient-echo echo-planar imaging (EPI) pulse sequence (TR = 1373 ms, TE = 33.4 ms, flip angle = 74°, 96 × 96 matrices, slice thickness = 2 mm, no gap, voxel size 2 mm × 2 mm × 2 mm, 72 axial slices covering the entire brain, 4 slices acquired at once). The first 5 volumes served as dummy scans and were discarded to avoid T1 equilibrium effects. Field maps were collected at the end of the experiment (TR = 400 ms, TE = 5.19 ms / 7.65 ms, flip angle = 60°, 64 × 64 matrices, slice thickness = 3 mm, 25% gap, resolution 3 mm isotropic, 32 axial slices). High-resolution anatomical T1-weighted images were acquired for each participant using a 3D MPRAGE sequence (192 axial slices, TR = 2250 ms, TI = 900 ms, TE = 2.99 ms, flip angle = 9°, field of view = 256 mm × 240 mm × 160 mm, matrix dimensions = 256 × 240 × 160, 1 mm isotropic resolution).

The data were preprocessed and analyzed using automatic analysis (aa) pipelines and modules (Cusack et al., 2015), which called relevant functions from Statistical Parametric Mapping software (SPM 12, http://www.fil.ion.ucl.ac.uk/spm) implemented in Matlab (The MathWorks, Inc., Natick, MA, USA). EPI images were realigned to correct for head motion using rigid-body transformation, unwarped based on the field maps to correct for voxel displacement due to magnetic-field inhomogeneity, and slice time corrected. The T1 image was coregistered to the mean EPI, and then coregistered and normalized to the MNI template. The normalization parameters of the T1 image were applied to all functional volumes. The data and models (see below) were temporally high-pass filtered with a cutoff at 1/128 Hz. Spatial smoothing of 10 mm FWHM was applied for the univariate whole-brain analysis, but not for the univariate ROI analysis or prior to multivariate analysis.

### Regions of interest (ROIs)

For the primary analysis, we focused on the MD and DMN networks (see Figure 4). The MD network was based on data from Fedorenko et al. (2013; Figure 2 in their paper), selecting frontoparietal regions responsive to cognitive demands across seven diverse tasks (http://imaging.mrc-cbu.cam.ac.uk/imaging/MDsystem). The DMN network was taken from Yeo et al. (2011), combining three subnetworks from the 17 network parcellation (numbers 15, 16, and 17) (Andrews-Hanna, 2012). The left and right hemispheres were averaged and projected back to both hemispheres to create a symmetrical volume (similar to Fedorenko et al., 2013). The combined networks were then smoothed at 4 mm FWHM to eliminate isolated voxels.

**Figure 2.**
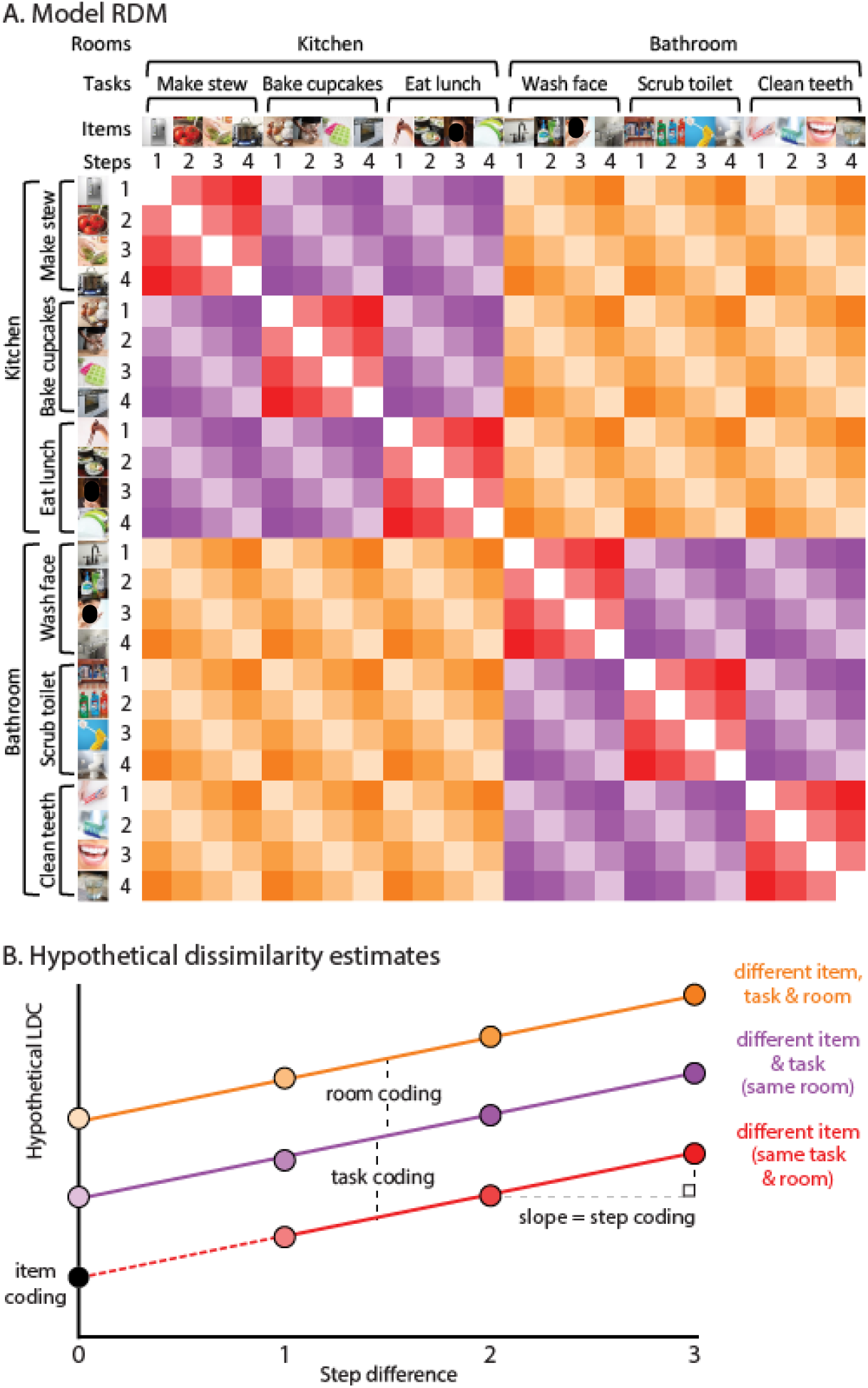
Illustration of representational similarity analysis. A. Simplified conceptual RDM. LDC dissimilarities are computed between every possible pair of events (6 cued tasks × 4 steps), generating a 24 × 24 RDM. Diagonal cells of the RDM are zero by definition as they do not reflect a dissimilarity between different events. Off-diagonal cells reflect pattern dissimilarity between events that always differ in search item, with varying additional differences in room, task, and step. B. Hypothetical pattern dissimilarities resulting from room, task, and item coding across step differences. Item coding can be estimated as the intercept, i.e., estimated LDC dissimilarity in in the absence of room, task, or step differences.

**Figure 3.**
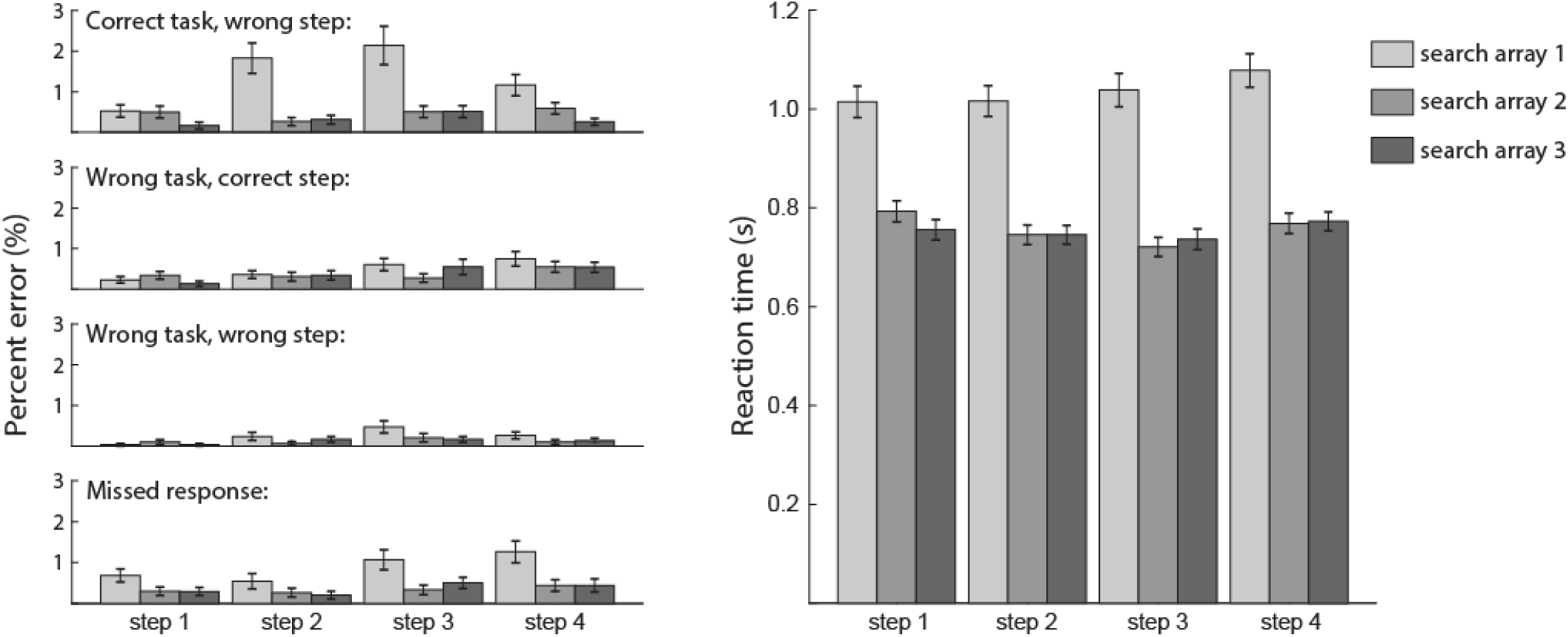
Behavioral performance summarized according to four possible error types (choosing an item from correct task but wrong step, wrong task but correct step, wrong task and step, and missed response), as well as reaction time for correct responses. For each step, the three bars indicate performance on each of the three successive search arrays. Error bars indicate standard error of the mean.

**Figure 4.**
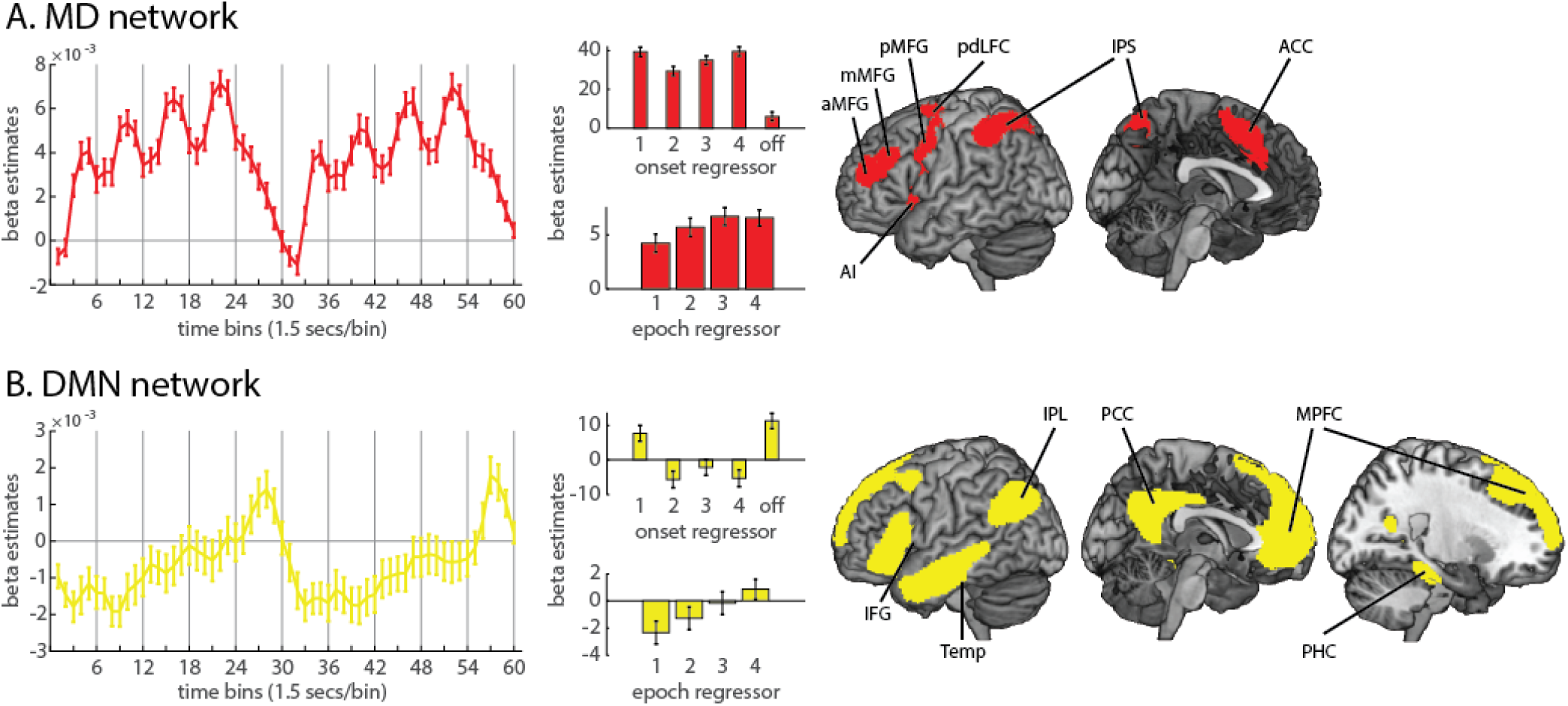
Univariate analysis of MD network (A) and DMN (B) BOLD response across task episodes. For each network, the left plot shows results of the FIR analysis, with BOLD response as a function of time as participants progressed across two consecutive episodes. The upper middle plot shows beta estimates associated with the step onsets (bars 1-4), and with the end of the episode (bar “off”). The lower middle plot shows beta estimates for epoch regressors for each step. Error bars indicate standard error of the mean. The right panel depicts component ROIs within each network.

Both the MD network (Dosenbach et al., 2006; Dosenbach et al., 2007; Crittenden et al., 2016) and the DMN (Andrews-Hanna et al., 2010; Andrews-Hanna, 2012; Wen et al., 2019) can be divided into finer components or subsystems, and following whole-network analysis, we examined separate sub-regions within each network. MD component ROIs (Mitchell et al., 2016) included three clusters along the anterior, middle, and posterior middle frontal gyrus (aMFG, mMFG, and pMFG), a posterior-dorsal region of lateral frontal cortex (pdLFC) in the superior precentral sulcus, and clusters in the intraparietal sulcus (IPS), anterior insula (AI), and anterior cingulate cortex (ACC). DMN component ROIs were defined as spatially separate clusters within the overall network, consisting of the medial prefrontal cortex (MPFC) and posterior cingulate cortex (PCC) along the midline, as well as the inferior frontal gyrus (IFG), inferior parietal lobule (IPL), parahippocampal cortex (PHC), and parts of the lateral temporal cortex extending to the temporal pole (Temp). Overlapping voxels of the AI and IFG were excluded from each ROI and their corresponding networks. Analyses were first performed using each network as a single large ROI, and then within each component ROI to examine more fine scale differences within each network. We controlled the False Discovery Rate (FDR) to correct for multiple comparisons across the number of networks (2) and component ROIs (13), respectively (Benjamini and Yekutieli, 2001).

### Univariate analysis

#### FIR Model

Statistical analyses were performed first at the individual level, using a general linear model (GLM). To capture the BOLD time-course throughout each task episode, as well as transitions between episodes, we modeled each consecutive pair of episodes. The first (dummy) episode was separately modeled and not analyzed. For the remaining data, a 90 s period starting from the onset of the first search array of every even number episode to the first search array of the next even number episode was modeled using a finite impulse response (FIR) basis set of 60 1.5 s boxcar regressors. In this way, the response throughout task episodes could be modelled without making assumptions about the shape of the hemodynamic response. Episodes with a high proportion of errors (episodes that had > 25% errors) were defined as error episodes, with the total number of error episodes per participant ranging from 0-6 (mean = 0.95, SD = 1.43). Any two consecutive error episodes were removed from the analysis using a similar but separate set of regressors. Effects of cues, and errors on individual search arrays, were also modeled separately, by convolving the duration of their respective events (1 s for cues and 2 s for error events) with a canonical hemodynamic response function. The six motion parameters and block means were included as regressors of no interest. Across the 90 s period, estimates for each FIR time bin were extracted from each whole network or component ROI, averaged over voxels within the region and across the six tasks. These average beta estimates for individual participants were entered into a random effects group analysis.

#### Event-based GLM analysis

To complement the FIR model, an event-based GLM analysis was performed. In this analysis, we aimed to separate phasic activity linked to onset of each step from tonic activity across the whole step within each episode. To control for the degree of visual difference between the search arrays of pairs of episodes, each combination of cued task × distractor task was modeled separately. For each combination, each step was modelled using two regressors, an onset regressor modelled with 0 s duration and an epoch regressor modelled with 9 s duration. Additionally, an offset regressor modelled with 0 s duration was placed at the end of the episode (following the final search array). Each regressor was convolved with the canonical hemodynamic response function. There were accordingly 162 regressors of interest, two (onset and epoch) for each of the four steps and one for the offset of the entire episode in each combination of six tasks × three possible distractor tasks from the other room (for example, the target task “make a stew” could be paired with distractor tasks “wash face”, “scrub toilet”, or “clean teeth”). Error episodes (defined as episodes that had > 25% errors) were removed from the analysis using a similar but separate set of regressors. The cue was modelled separately using a similar combination of onset (0 s duration) and epoch (duration from cue onset to the onset of the first task step) regressors. Motion parameters and block mean regressors were included as before. Beta estimates were averaged across the 18 cued task × distractor task combinations for individual participants, and entered into random effects group analyses. We first examined the mean effect of onset/offset and epoch regressors versus implicit baseline (with FDR correction across ROIs). Repeated measures analyses of variance (ANOVAs) were used to examine changes across steps, including linear and quadratic trends. To complement the ROI analyses, contrasts were also carried out at the whole brain level, using a voxel-wise FDR-corrected threshold of p < 0.025 per tail. Results were rendered using MRIcroGL (version 1.2.20190902; www.nitrc.org/projects/mricrogl).

### RSA analysis

We performed representational similarity analysis (RSA) using the linear discriminant contrast (LDC) to quantify dissimilarities between activation patterns. The analysis used the RSA toolbox (Nili et al., 2014), in conjunction with in-house software. The LDC was chosen because it is multivariate noise-normalized, potentially increasing sensitivity, and is a cross-validated measure which is distributed around zero when the true distance is zero (Walther et al., 2016). The LDC also allows inference on contrasts of dissimilarities across multiple pairs of task events. A pattern for each step of each combination of cued task and distractor task was obtained, by averaging the onset and epoched responses from the event-based GLM described above. This resulted in 72 patterns in total in each run. For each pair of patterns, the patterns from run 1 were projected onto a Fisher discriminant fitted for run 2, with the difference between the projected patterns providing a cross-validated estimate of a squared Mahalanobis distance. This was repeated projecting run 2 onto run 1, and we took the average as the dissimilarity measure between the two patterns. All pairs of pattern dissimilarities therefore formed a symmetrical representational dissimilarity matrix (RDM) with zeros on the diagonal by definition. To allow comparison of dissimilarity magnitude across ROIs of different sizes, the LDC values were normalized by dividing by the number of voxels within each ROI.

As for univariate analyses, we first performed RSA analysis using activation patterns from the DMN and MD networks treated as single large ROIs, and then repeated it on component ROIs. To introduce measures for room, task, step and item coding, Figure 2A shows a simplified version of the full 72 × 72 representational dissimilarity matrix (RDM), collapsing across distractor task to produce just a 24 × 24 matrix. In this matrix, each cell represents a cross-validated LDC dissimilarity between the corresponding two task events. These included event pairs that shared the same cued task (red cells; e.g., “take food from fridge” and “wash vegetables”); events that shared the same room but different cued tasks (purple cells; e.g., “take food from fridge” and “hand mix batter”); and events that differed in both cued task and room (orange cells; e.g. “take food from fridge” and “use facial wash”). All event pairs additionally differed in item. Saturation of the colors is used to indicate the difference in steps between event pairs. The cells on the diagonal (white) are zero by definition as they do not reflect a comparison between different task events.

To extract measures for room, task, step and item coding, we fit values in the matrix with the regression model illustrated in Figure 2B. In this model, the LDC estimate for any entry in the matrix is the linear sum of components from differences in target item (contributing equally to all cells; major diagonal of the matrix ignored), room, task, and step. As a measure of step coding, we used the slope of the function relating LDC estimate to step difference. For example, steps 1 and 2 have a step difference of 1, while steps 1 and 4 have a step difference of 3. As a measure of room coding, we used the difference in LDC estimate for different room and same room/different task cases (Figure 2B, orange vs. purple). As a measure of task coding, we used the difference in LDC dissimilarity for same room/different task and same task cases (Figure 2B, purple vs red). As item difference contributed similarly to all cells, it was estimated as the intercept of the full model (Figure 2B, black dot).

For the actual fitting, we used a more complex model based on the full 72 × 72 RDM, used to remove a potential visual confound (see Extended Data Figure 2-1). For any cell in the full RDM, search arrays could share items from zero, one, or two tasks. For example, consider an episode with cued task “make a stew” and distractor items coming from the distractor task “wash face”. Search arrays from this episode would share no items with search arrays from an episode of “bake cupcakes” with distractors from “scrub toilet”; arrays would share items from one task when compared to the episode “make a stew” with distractors from “wash face”; arrays would share items from both tasks when compared to the episode “wash face” with distractors from “make a stew”. In the full model, we added an additional regressor to remove this potential visual confound. This was defined as “visual difference”, with values of 1 for no shared tasks, 0.5 for one shared task, and 0 for two shared tasks. The mean coding estimates across subjects were tested against zero using 1-tailed t-tests, and multiple comparisons across ROIs were corrected using FDR <0.05 per measure.

### Experimental Design and Statistical Analysis

All statistical tests were performed across 42 participants (20 male, 22 female), with no between-subject factors. Behavioral analyses used repeated measures ANOVA to compare conditions. Univariate fMRI analyses used one-sample (paired–sample) two-tailed t-tests to compare responses against baseline, between conditions, or linear contrasts of regression coefficients, and repeated measures ANOVA to compare multiple conditions. RSA fMRI analyses used one-sample one-tailed t-tests to test for greater-than-chance encoding of each information type, paired-sample two-tailed t-tests to compare networks, and repeated measures ANOVA to test the interaction of information type and network. Within-subject factors are detailed in the relevant Results sections. For each analysis, multiple comparisons (across 2 networks, 13 component ROIs, or all brain voxels) were accounted for by controlling the FDR at 0.05, unless noted otherwise. Effect sizes were calculated using partial Eta-squared for ANOVAs and Cohen’s d for t-tests. Analyses were performed using Matlab (The MathWorks, Inc., Natick, MA, USA), SPM 12 (http://www.fil.ion.ucl.ac.uk/spm), and SPSS (version 25). In repeated measures ANOVA, Greenhouse-Geisser correction was used to adjust for non-sphericity. Data are available on request.

## Results

### Behavioral results

Group behavioral performance is shown in Figure 3. Error rates were calculated after removal of occasional whole error episodes (defined as episodes that had > 25% errors, suggesting that the correct cue was not being followed; 0-6 across participants), and reaction time was calculated for only correct trials. Overall accuracy was 97.5% ± 0.4% (mean ± SEM) and overall reaction time was 849 ± 23 ms. Error responses were broken into four error types: choosing an item from the correct task but wrong step, wrong task but correct step, wrong task and step, and missed response. Results show poorest performance for the first search array of each step, when participants were required to switch from one step to the next. A step (steps 1-4) × search array (first, second, third within each step) ANOVA was performed for each type of error. All error types showed a main effect of step (all F(3,123) > 3.79, all ps < 0.03, all η_p_^2^ > 0.08), and linear trend analyses indicated an overall increase in error across steps (all F(1,41) > 7.70, all ps < 0.01, all η_p_^2^ > 0.15). A main effect of search array was found for correct task wrong step errors, as well as for missed responses (both F(2,82) > 14.36; both ps < 0.001, both η_p_^2^ > 0.25), reflecting higher errors on the first search array of each step. Finally, correct task wrong step errors showed a significant step × array interaction (F(6,246) = 5.96, p < 0.001, η_p_^2^ = 0.13). A similar ANOVA for reaction time also showed a significant main effect for step (F(3,123) = 17.21, p < 0.001, η_p_^2^ = 0.30), a significant main effect for search array (F(2,82) = 234.42, p < 0.001, η_p_^2^ = 0.85), and a significant step × array interaction (F(6,246) = 9.83, p < 0.001, η_p_^2^ = 0.19).

### Univariate results

#### ROI analysis

The FIR model provided estimates of the observed BOLD response time-course across a pair of task episodes, in successive 1.5 s windows starting from the onset of the first step. In the main analysis, we extracted these FIR responses from *a priori* networks (Figure 4, left). The MD network exhibited positive activity throughout each episode, along with four peaks corresponding to the four steps. These results suggest involvement in setting up and executing individual task steps. Additionally, overall MD activity gradually increased throughout the task episode, suggesting that the MD network is also sensitive to progress through the episode. For DMN regions, in contrast, tonic activation began below baseline but gradually increased through the episode, culminating in a large phasic response at episode completion. For both networks, the signal clearly resets between episodes.

To quantify the phasic and tonic components contributing to the BOLD response at each task step, we performed a complementary GLM analysis with onset and epoch regressors modelling each task step (Figure 4, middle). Four onset regressors were designed to reflect phasic activity at the onset of each task step. The final offset regressor was included to capture the phasic activity at the end of the episode. Finally, epoch regressors were designed to reflect tonic activity throughout each step.

Within the MD network, there were strong onset responses, in line with FIR results. Contrasts with baseline showed that all four step onsets were significantly greater than baseline (all *t*s > 10.91, all *p*s < 0.001, all ds > 1.68) and there was a smaller yet significant offset response (t = 2.48, p = 0.02, d = 0.38). A one-way repeated measures ANOVA showed a significant difference across the four step onsets (F(3,123) = 5.60, p < 0.01, η_p_^2^ = 0.12), with a quadratic (F(1,41) = 21.61, p < 0.001, η_p_^2^ = 0.35) but not linear (F(1,41) = 0.22, p = 0.64, η_p_^2^ < 0.01) trend across steps, reflecting an increasing response across steps 2-4, but a disproportionate response to the onset of the first step, i.e. the onset of the entire episode. Looking at epoch regressors, all four epoch responses were greater than baseline (all *t*s > 3.96, all *ps* < 0.001, all ds > 0.61). ANOVA showed a significant main effect of step (F(3,123) = 7.73, p = 0.01, η_p_^2^ = 0.16), as well as a significant linear (F(1,41) = 9.48, p < 0.01, η_p_^2^ = 0.19) and quadratic trend (F(1,41) = 5.08, p = 0.03, η_p_^2^ = 0.11), reflecting an increasing but saturating response.

The DMN network showed a different profile. Only the onset of the first step (*t* = 3.22, *p* < 0.01, d = 0.50) and the offset response at the end of the episode (*t* = 4.38, *p* < 0.001, d = 0.68) were greater than baseline. Step onsets 2-4 were not significantly different from baseline (all |t|s < 2.09, all ps > 0.07, all |d|s < 0.33). ANOVA of the four step onsets showed a significant main effect of step (F(3,123) = 9.87, p < 0.001, η_p_^2^ = 0.19), as well as significant linear (F(1,41) = 9.70, p < 0.01, η_p_^2^ = 0.19) and quadratic (F(1,41) = 7.16, p = 0.01, η_p_^2^ = 0.15) trends, consistent with the larger response to the first onset. Among the epoch responses, the first step was significantly lower than baseline (t = −3.21, p = 0.01, d = −0.49; for steps 2-4 all |t|s < 1.60, all ps > 0.23, all |d|s < 0.19). ANOVA showed a significant main effect of step (F(3,123) = 18.42, p < 0.001, η_p_^2^ = 0.31), as well as a significant linear trend (F(1,41) = 38.89, p < 0.001, η_p_^2^ = 0.49), suggesting an increase in activation across steps. As seen in the FIR time-course, this implies a gradual release of tonic deactivation across the duration of the task episode.

To examine whether the profiles of different regions within each network showed unique responses, we performed the same analyses (FIR time-course modelling and event-based GLM) on individual ROIs (see Extended Data Figure 4-1). Trends of activation across the four steps for individual ROIs were largely similar to the network in which they belong, though there were some differences between ROIs. Within the MD network, aMFG and AI showed negative epoch responses, in contrast to other regions. The episode offset response was also especially high in aMPFC and especially small in pdLFC and IPS. Within the DMN, PHC showed positive epoch responses, in contrast to other regions.

#### Whole-brain analysis

Results from the whole-brain analysis, again separating onset and epoch regressors, are shown in Figure 5. Panel A shows responses within an episode, with the left-hand side showing contrasts of the mean onset (i) and epoch (ii) response against baseline, and the right-hand side showing increasing trends across steps for the onset response (iii) and the epoch response (iv). The onset of step 1 and the offset of step 4 are special, since these correspond to the onset and offset of a whole episode, and it is evident from the ROI analysis that their neural response also differs from step onsets within an episode. Therefore, these episode-boundary responses were not included in the within-episode contrasts, but were instead examined separately. Panel B of figure 5 shows onset responses at episode onset (left) and offset (right), contrasted against both baseline (i, iii) and against the adjacent step onset response (ii, iv).

**Figure 5.**
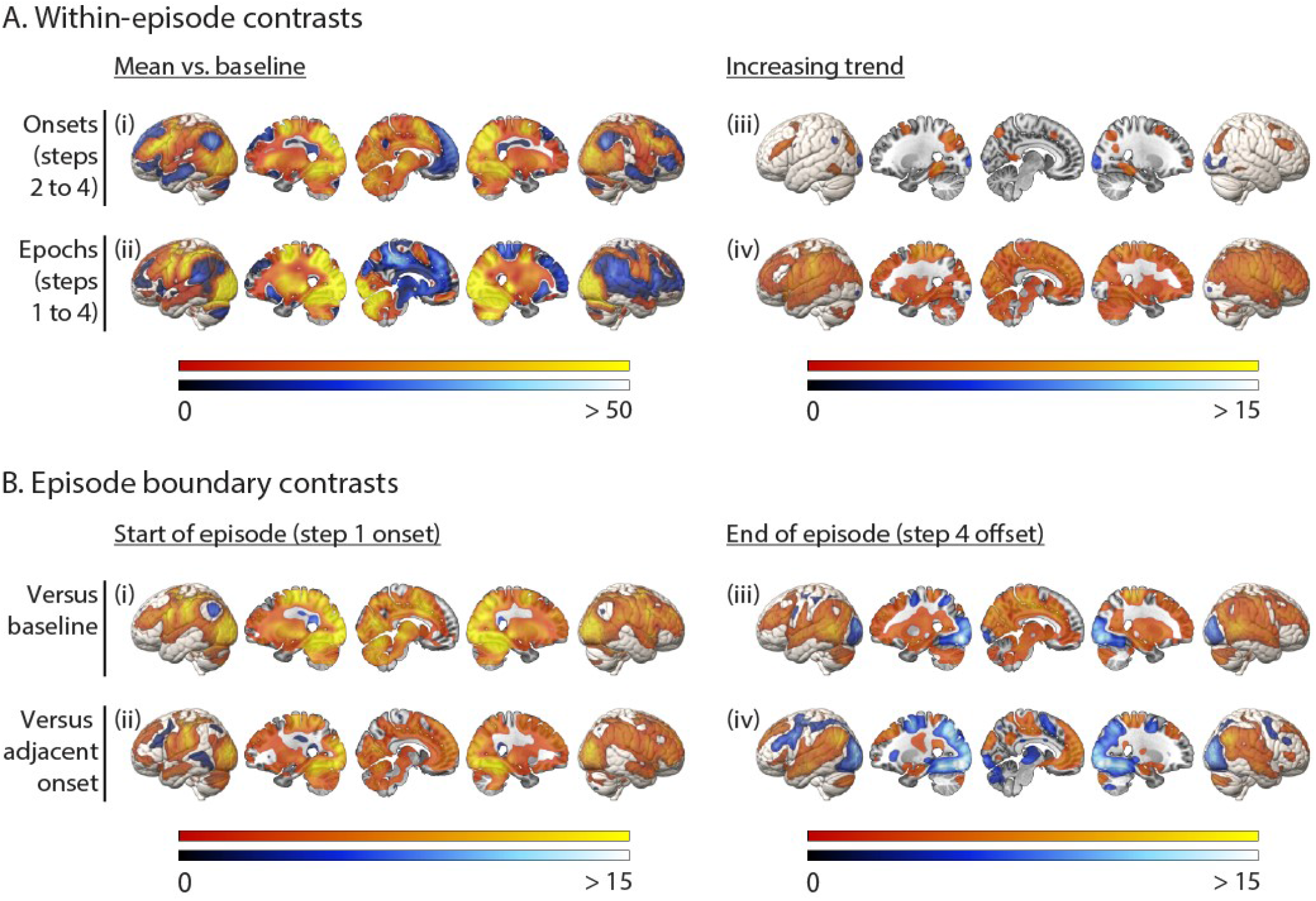
Whole brain univariate analysis. (A) Responses within an episode, including (i) mean phasic responses to the onset of each step; (ii) mean tonic responses across the duration of each step; (iii) increases in the phasic response across step onsets; (iv) increases in the tonic response across step epochs. (B) Transient responses at episode boundaries, including (i) episode onset versus baseline; (ii) episode onset versus step 2 onset; (iii) episode offset versus baseline; (iv) episode offset versus step 4 onset. Colors indicate t-values, with warm and cool scales indicating positive and negative tails respectively. All activation maps are thresholded at FDR < 0.025 per tail.

In comparison to baseline, the mean step onset response (Figure 5Ai) was significantly positive throughout the MD network, as well as visual cortex, motor cortex, and subcortical structures including the cerebellum. The mean step onset response was significantly negative throughout the DMN. Mean epoch responses greater than baseline (Figure 5Aii) were also extensive, including parietal and frontal regions overlapping with the MD ROIs, as well as expected regions of visual and motor cortex. Again, we saw negative epoch responses in much of the DMN. We next examined activity changes across steps within an episode. An increase in the amplitude of the step onset response was restricted to MD regions (Figure 5Aiii). In contrast, a linear increase in the tonic epoch response was widespread across most of the brain (Figure 5Aiv). The only exception was areas of visual cortex, where both onset and epoch responses decreased across an episode. Finally, we were interested in the response at episode boundaries, i.e. the onset of the first step (initiation of an episode) and the offset of the fourth step (completion of an episode). The response to step 1 onset was substantial across much of the brain, whether compared to baseline or to step 2 onset (Figure 5Bi and 5Bii), including visual cortex and parts of DMN and MD networks. The episode completion response was also significantly greater than baseline in many brain regions (Figure 5Biii), including parts of both MD and DMN networks, while deactivations were mainly observed in visual cortex. Interestingly, this response exceeded the previous step onset response in the DMN but not the MD network (Figure 5Biv).

The results may be summarized as follows. Most MD regions, along with visual cortex, showed positive onset and epoch responses to all steps, suggesting direct involvement in setting up and executing task steps. Sensitivity to the large-scale structure of the task episode was evident in gradually increasing activity as the episode progressed, along with phasic responses at onset and offset of the whole episode. Interestingly, tonic ramping activity was widespread throughout the brain, whereas increasing onset responses were highly specific to the MD network, and an episode offset response exceeding the preceding step onset was largely specific to the DMN.

### RSA results

Results of the RSA analysis are shown in Figure 6. In panel A, LDC representational distance estimates are plotted for various comparisons of events (see also Figure 2), based on activation patterns across the DMN and MD networks. The coefficients of the linear model fit to these data are plotted in panel B, quantifying coding of different types of information – room, cued task, step, and item. Analyses of individual ROIs within the two networks are shown in Extended Data Figure 6-1.

**Figure 6.**
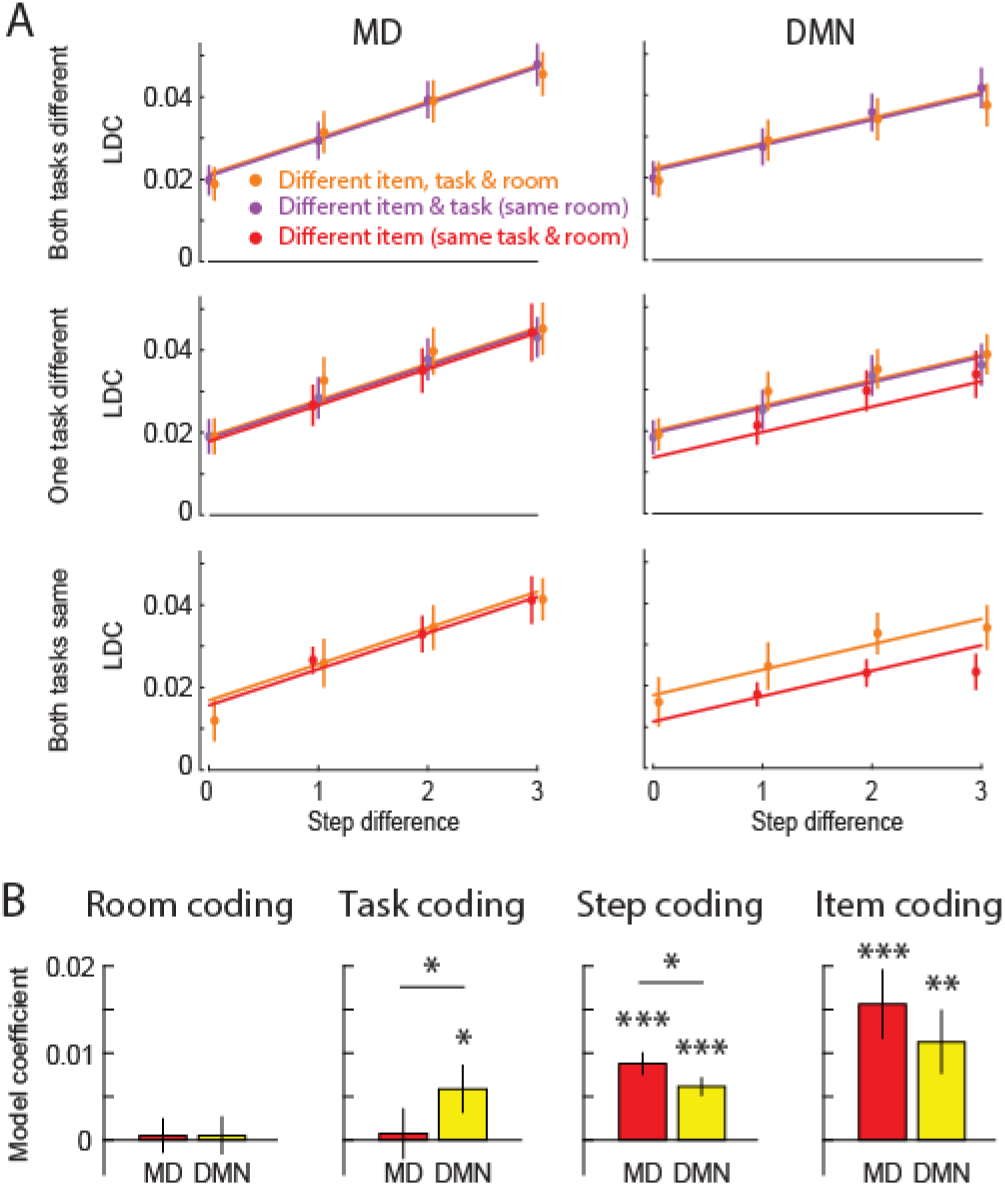
(A) LDC representational distance estimates, modelled as in figure 2b. To avoid clutter, data points are plotted separately for different levels of visual difference (0%, 50%, 100% task overlap), and are offset fractionally along the x-axis. Lines represent the mean linear model fit, estimated using all data (except matched-item comparisons). (B) LDC contrasts representing strength of room, cued task, step, and item coding in DMN and MD network-level ROIs. Asterisks above each bar indicate significance of 1-tailed t-tests against zero, after controlling FDR <0.05 across ROIs; horizontal lines indicate a significant 2-tailed paired t-test between networks; *** p < 0.001, ** p < 0.01, * p < 0.05. Error bars represent +/− 1 standard error of the mean across subjects.

#### Network comparison

First, we asked whether activity patterns in the MD and DMN networks differentially carried information about distinct aspects of task episodes. A 2 (network) × 4 (type of information) repeated measures ANOVA showed a significant interaction (F(4,164) = 3.13, p = 0.033), as well as a main effect of information type (F(4,164) = 3.65, p = 0.016) but not network (F(1,41) = 0.25, p = 0.62). The interaction was driven by the DMN having a relative preference for coding the identity of the cued task, while the MD network had a relative preference for step-level coding (step position, and item identity). We next assessed coding of each information type in turn.

#### Room coding

Neither the DMN nor MD network ROIs showed significant room coding (both *t*s < 0.25, both *p*s > 0.39, both *d*s < 0.04). Similarly, none of the individual ROIs showed significant room coding (all *t*s < 1.70, all *p*s > 0.11, *d*s < 0.27), though it was numerically strongest in the MPFC.

#### Task coding

The DMN network showed significant coding of the cued task (t = 2.18, p = 0.02, d = 0.33), while the MD network did not (t = 0.26, p = 0.40, d = 0.04); the difference between networks was also significant (t = 2.52, p = 0.02, d = 0.39). None of the individual ROIs showed significant task coding after FDR correction for multiple comparisons across ROIs. PCC, and ACC showed task coding before correction (both ts > 1.74, both ps < 0.044, both ds > 0.27). Task coding was positive in all six DMN ROIs, but only four of seven MD ROIs.

It is possible that the response to regressors modelling adjacent steps could be similar due to imperfect temporal separation of the signal, such that pairs of steps within the same task appear more similar than those from different tasks due to differences in temporal separation in addition to differences in task identity. We examined this possibility by fitting four separate linear regression models using subsets of cells, chosen to differ in separation of one, two, or three steps. That is, we extracted LDC values from cells of the DMN network RDM that represented one step apart (1 vs. 2, 2 vs. 3, and 3 vs. 4), two steps apart (1 vs. 3 and 2 vs. 4), or three steps apart (1 vs. 4), and, in each case, fitted a model with room, cued task, and visual difference regressors. If temporal proximity were contributing to activity pattern similarity, and hence to apparent task coding in the DMN, we should expect a stronger effect for steps closer together in time. However, we found no evidence of any difference in task coding across these three conditions (F(2, 82) = 0.39, p = 0.61, η_p_^2^ = 0.01), nor a linear trend as a function of step (F(1,41) = 0.44, p = 0.51, η_p_^2^ = 0.01). Task coding as a function of step difference is shown in Extended Data Figure 6-2.

#### Step coding

Step coding was significant in both the DMN (t = 6.34, p < 0.001, d = 0.98) and MD (t = 7.25, p < 0.001, d = 1.12) network ROIs. The MD network showed significantly greater step coding than the DMN (t = 2.38, p =0.02, d = 0.37). Step coding was also significant in all the individual ROIs (all ts > 2.02, all ps < 0.03, all ds > 0.31). This was not surprising, as in our univariate analysis we observed strong linear trends across the task episode for most of the brain (see Figure 5Aiv).

#### Item coding

Both DMN (t = 3.15, p = 0.002, d = 0.49) and MD (t = 4.00, p < 0.001, d = 0.62) network ROIs showed significant coding of item. The two networks did not significantly differ in item coding (t = 1.88, p = 0.07, d = 0.29). In the individual ROIs, item coding was especially strong in parietal regions, with only IPS (t = 4.58, p < 0.001, d = 0.71) and IPL (t = 2.89, p < 0.01, d = 0.44) showing significant item coding after FDR correction for multiple comparisons (prior to correction, item coding was also present in ACC (t = 2.05, p = 0.02, d = 0.32) and PCC (t = 1.90, p = 0.03, d = 0.29)).

## Discussion

We used fMRI to examine how cortical networks respond to and represent different aspects of multi-step task episodes. Specifically, we focused on the DMN and MD networks. Using FIR and event-related analyses to capture the evolution of the BOLD response throughout a multi-step episode, we found that MD regions showed positive activity throughout the episode, with separate peaks for successive steps. These results suggest involvement in setting up and executing individual task steps. In contrast, the DMN showed overall deactivation. Along with widespread other regions, both DMN and MD showed sensitivity to the large-scale structure of the episode, with phasic responses to onset and offset, and gradually increasing tonic activity as the episode progressed. MD regions were unique in showing an increase in the phasic step response across the episode, while a characteristic feature of DMN regions was an episode offset response that exceeded the final step response.

Representational similarity analysis indicated that MD regions showed strong coding of individual items but not the identity of the cued task, while for DMN, both item and task coding were significant. Both networks additionally showed strong step coding, which was significantly greater in the MD network than the DMN. The finding that MD regions are especially sensitive to the identity of a current task step and its specific item content is consistent with prior research. Many previous experiments have shown coding of task-relevant information in MD regions that can rapidly change according to task demands (Freedman et al., 2001; Li et al., 2007; Woolgar et al., 2011), including radical reorganization between successive task steps (Sigala et al., 2008). fMRI studies show strong MD activity when a sub-goal is completed and in transitions from one event to another (Sridharan et al., 2007; Farooqui et al., 2012), with progressively increasing activity as a goal is approached (Farooqui et al., 2012; Desrochers et al., 2015; Desrochers et al., 2019). The pattern of MD activation in our study is broadly consistent with these previous findings. The results suggest that, as a task episode progresses, MD representations in particular are in constant flux, reorganizing to encode the detailed contents of each task step. Representational content includes the position of the step within the task episode and the identity of the item required on that step, which, in the current task, may serve as an attentional template for visual search decisions (Desimone and Duncan, 1995).

Within the DMN, univariate results showed a burst of activity at the beginning of each episode, but no significant onset responses to subsequent steps. At the completion of the episode, DMN also exhibited an especially strong offset response. These findings are consistent with prior reports of transient DMN activation at event boundaries (Speer et al., 2007), including transitions to a new task (Smith et al., 2018). Our data showed, however, that both onset and offset responses were widespread in the brain (Fox et al., 2005) and it was the magnitude of the offset response that was most specific to the DMN. It has been proposed that the mental programs required for carrying out a task are assembled at the beginning of task execution (Schneider and Logan, 2006; Farooqui and Manly, 2019). It is possible that the DMN, along with multiple other brain regions, is involved in long-term memory retrieval for the entire task sequence at episode initiation, and long-term memory storage at episode completion. The DMN’s marked response to task episode boundaries, along with its significant representation of task identity, are consistent with the proposal that the DMN has long temporal receptive windows and codes for information accumulated over long time scales (Hasson et al., 2008; Lerner et al., 2011; Chen et al., 2016).

For the DMN as a whole, significant coding of cued task identity suggests representation of broad task context. A particular role of the DMN in task coding is consistent with its proposed function in representing “situation models” — higher-level cognitive representations of relationships between different elements of an episode — as part of a “posterior medial system” centered on PHC and retrosplenial cortex (Diana et al., 2007; Ranganath and Ritchey, 2012; Reagh and Ranganath, 2018). The MPFC sub-region of the DMN has been implicated in schema representation, capturing similarities across multiple episodes (Preston and Eichenbaum, 2013; Ghosh and Gilboa, 2014; Robin and Moscovitch, 2017), and might therefore have been expected to show room coding. Although room coding was numerically strongest in this ROI, it was not significant. Stronger room coding might have been observed if the grouping by room had been behaviorally relevant rather than incidental.

We also found that the DMN showed significant coding for items, and thus represents not just full tasks, but also specific contents within a task episode. Although we found both item and task representations coexisting in the DMN, consistent with a compositional code, this experiment cannot determine whether and where items and tasks might be bound into a conjunctive representation, or maintained as independent factorized components (Behrens et al., 2018): because items were unique to each task, item-task conjunctions are indistinguishable from item coding. Disentangling these different forms of co-representation requires designs where the same item appears in different contexts. Such designs have identified item-context conjunctions in the hippocampus (Hsieh et al., 2014) item-order associations in various frontal and temporal regions (Reverberi et al., 2012; Kalm and Norris, 2014), rule-rule compositionality in lateral frontal cortex (Cole et al., 2011), and factorized coding of sequence and position in midcingulate cortex (Holroyd et al., 2018) and in electrophysiological signals during learning and replay (Liu et al., 2019).

Both MD and DMN, along with most regions of the brain, tracked progress through the task episode, shown by increasing trends in the univariate data. These observations are consistent with previous studies that reported ramping activity across task episodes in specific MD (Farooqui et al., 2012; Desrochers et al., 2015; Desrochers et al., 2019) and DMN regions (Farooqui and Manly, 2018), but suggest that it might be a much more global property of brain function (Farooqui and Manly, 2018). While parts of visual cortex showed a decrease in sustained activity over the episode, which may reflect adaptation to the sensory input (Grill-Spector and Malach, 2001; Grill-Spector et al., 2006), most other cortical regions showed an increase in sustained activity over the episode, which reset at episode boundaries. As this effect was so widespread, it is difficult to offer a precise interpretation, and different areas may increase for different reasons (Kalm and Norris, 2017). For example, it is possible that increased activations in some regions reflect revision and reconfiguration of control representations that may increase in demand as larger portions of the task are complete (Farooqui et al., 2012; Desrochers et al., 2015; Desrochers et al., 2019), or expectation of, and preparation for, trial completion (Shidara and Richmond, 2002). These activity changes could also reflect gradual accumulation and integration of new information into an episode representation (Hasson et al., 2008; Dumontheil et al., 2011; Lerner et al., 2011). The global nature of the signal is potentially consistent with models of multi-step decision-making, in which evidence accumulation is massively parallel within temporal chunks that are serially chained (Zylberberg et al., 2011; Dehaene and Sigman, 2012). In rats, anticipation of distant goals has been associated with slowly ramping dopamine release (Howe et al., 2013), which could suggest a mechanism for the widespread cortical effects observed here. In contrast to the global nature of the tonically increasing response, increases in the phasic step response appeared highly specific to the MD network. Speculatively, the phasic MD response may track progress in discrete steps, in addition to the tonic global signal which reflects a more continuous measure of progress. A similar distinction between neural signals that track progress in a smooth versus action-linked manner has also been observed in the rat brain (Ma et al., 2014).

A hierarchical control structure is an organized representation of control elements, including task identity, local entities, and serial position codes (Rosenbaum et al., 1983; Schneider and Logan, 2006). Our results describe how broad brain networks are involved in the execution of task sequences, with MD and DMN regions exhibiting distinct time-courses throughout the episode, and different profiles of information representation. The DMN, we suggest, may establish overall cognitive context, representing both individual cognitive operations and their broader context, and perhaps involved in binding them together. These functions may be consistent with representing a “situation model” (Ranganath and Ritchey, 2012). At the same time, the MD system, along with sensory regions, processes and tracks the detailed content of individual cognitive operations, locked to discrete events within the episode. Both networks respond to the broad temporal structure of task episodes, with phasic activity at task onset and offset, and gradually increasing activity as the whole sequence of steps progresses. Acting together, they reflect the hierarchical structure of goal-directed behavior.

## Acknowledgements

This work was supported by funding from the Medical Research Council (United Kingdom), program SUAG/045/G101400. TW was supported by the Medical Research Council PhD Studentship, Taiwan Cambridge Scholarship from the Cambridge Commonwealth, European & International Trust, and the Percy Lander studentship from Downing College.

## References

Andrews-Hanna JR (2012) The brain’s default network and its adaptive role in internal mentation. Neuroscientist 18:251–270.

Andrews-Hanna JR, Reidler JS, Sepulcre J, Poulin R, Buckner RL (2010) Functional-anatomic fractionation of the brain’s default network. Neuron 65:550–562.

Asaad WF, Rainer G, Miller EK (2000) Task-specific neural activity in the primate prefrontal cortex. J Neurophysiol 84:451–459.

Baldassano C, Hasson U, Norman KA (2018) Representation of Real-World Event Schemas during Narrative Perception. J Neurosci 38:9689–9699.

Behrens TEJ, Muller TH, Whittington JCR, Mark S, Baram AB, Stachenfeld KL, Kurth-Nelson Z (2018) What Is a Cognitive Map? Organizing Knowledge for Flexible Behavior. Neuron 100:490–509.

Ben-Yakov A, Henson RN (2018) The Hippocampal Film Editor: Sensitivity and Specificity to Event Boundaries in Continuous Experience. J Neurosci 38:10057–10068.

Ben-Yakov A, Eshel N, Dudai Y (2013) Hippocampal immediate poststimulus activity in the encoding of consecutive naturalistic episodes. J Exp Psychol Gen 142:1255–1263.

Benjamini Y, Yekutieli D (2001) The control of the false discovery rate in multiple testing under dependency. Annals of Statistics 29:1165–1188.

Chen J, Honey CJ, Simony E, Arcaro MJ, Norman KA, Hasson U (2016) Accessing Real-Life Episodic Information from Minutes versus Hours Earlier Modulates Hippocampal and High-Order Cortical Dynamics. Cereb Cortex 26:3428–3441.

Cohn-Sheehy BI, Ranganath C (2017) Time Regained: How the Human Brain Constructs Memory for Time. Curr Opin Behav Sci 17:169–177.

Cole MW, Etzel JA, Zacks JM, Schneider W, Braver TS (2011) Rapid transfer of abstract rules to novel contexts in human lateral prefrontal cortex. Front Hum Neurosci 5:142.

Cooper R, Shallice T (2000) Contention scheduling and the control of routine activities. Cogn Neuropsychol 17:297–338.

Crittenden BM, Mitchell DJ, Duncan J (2015) Recruitment of the default mode network during a demanding act of executive control. Elife 4:e06481.

Crittenden BM, Mitchell DJ, Duncan J (2016) Task Encoding across the Multiple Demand Cortex Is Consistent with a Frontoparietal and Cingulo-Opercular Dual Networks Distinction. J Neurosci 36:6147–6155.

Cusack R, Vicente-Grabovetsky A, Mitchell DJ, Wild C, Auer T, Linke AC, Peelle JE (2015) Automatic analysis (aa): Efficient neuroimaging workflows and parallel processing using Matlab and XML. Frontiers in Neuroinformatics 8.

Dale AM (1999) Optimal experimental design for event-related fMRI. Hum Brain Mapp 8:109–114.

Dehaene S, Sigman M (2012) From a single decision to a multi-step algorithm. Curr Opin Neurobiol 22:937–945.

Desimone R, Duncan J (1995) Neural mechanisms of selective visual attention. Annu Rev Neurosci 18:193–222.

Desrochers TM, Chatham CH, Badre D (2015) The Necessity of Rostrolateral Prefrontal Cortex for Higher-Level Sequential Behavior. Neuron 87:1357–1368.

Desrochers TM, Collins AGE, Badre D (2019) Sequential Control Underlies Robust Ramping Dynamics in the Rostrolateral Prefrontal Cortex. J Neurosci 39:1471–1483.

Diana RA, Yonelinas AP, Ranganath C (2007) Imaging recollection and familiarity in the medial temporal lobe: a three-component model. Trends Cogn Sci 11:379–386.

Dosenbach NU, Visscher KM, Palmer ED, Miezin FM, Wenger KK, Kang HC, Burgund ED, Grimes AL, Schlaggar BL, Petersen SE (2006) A core system for the implementation of task sets. Neuron 50:799–812.

Dosenbach NU, Fair DA, Miezin FM, Cohen AL, Wenger KK, Dosenbach RA, Fox MD, Snyder AZ, Vincent JL, Raichle ME, Schlaggar BL, Petersen SE (2007) Distinct brain networks for adaptive and stable task control in humans. Proc Natl Acad Sci U S A 104:11073–11078.

Dumontheil I, Thompson R, Duncan J (2011) Assembly and use of new task rules in fronto-parietal cortex. J Cogn Neurosci 23:168–182.

Duncan J (2010) The multiple-demand (MD) system of the primate brain: mental programs for intelligent behaviour. Trends Cogn Sci 14:172–179.

Duncan J (2013) The structure of cognition: attentional episodes in mind and brain. Neuron 80:35–50.

Everling S, Tinsley CJ, Gaffan D, Duncan J (2002) Filtering of neural signals by focused attention in the monkey prefrontal cortex. Nat Neurosci 5:671–676.

Ezzyat Y, Davachi L (2011) What constitutes an episode in episodic memory? Psychol Sci 22:243–252.

Farooqui AA, Manly T (2018) Hierarchical Cognition Causes Task-Related Deactivations but Not Just in Default Mode Regions. eNeuro 5.

Farooqui AA, Manly T (2019) We do as we construe: extended behavior construed as one task is executed as one cognitive entity. Psychological Research-Psychologische Forschung 83:84–103.

Farooqui AA, Mitchell D, Thompson R, Duncan J (2012) Hierarchical organization of cognition reflected in distributed frontoparietal activity. J Neurosci 32:17373–17381.

Fedorenko E, Duncan J, Kanwisher N (2013) Broad domain generality in focal regions of frontal and parietal cortex. Proc Natl Acad Sci U S A 110:16616–16621.

Fox MD, Snyder AZ, Barch DM, Gusnard DA, Raichle ME (2005) Transient BOLD responses at block transitions. Neuroimage 28:956–966.

Freedman DJ, Riesenhuber M, Poggio T, Miller EK (2001) Categorical representation of visual stimuli in the primate prefrontal cortex. Science 291:312–316.

Ghosh VE, Gilboa A (2014) What is a memory schema? A historical perspective on current neuroscience literature. Neuropsychologia 53:104–114.

Grill-Spector K, Malach R (2001) fMR-adaptation: a tool for studying the functional properties of human cortical neurons. Acta Psychol (Amst) 107:293–321.

Grill-Spector K, Henson R, Martin A (2006) Repetition and the brain: neural models of stimulus-specific effects. Trends Cogn Sci 10:14–23.

Hasson U, Yang E, Vallines I, Heeger DJ, Rubin N (2008) A hierarchy of temporal receptive windows in human cortex. J Neurosci 28:2539–2550.

Holroyd CB, Ribas-Fernandes JJF, Shahnazian D, Silvetti M, Verguts T (2018) Human midcingulate cortex encodes distributed representations of task progress. Proc Natl Acad Sci U S A 115:6398–6403.

Howe MW, Tierney PL, Sandberg SG, Phillips PE, Graybiel AM (2013) Prolonged dopamine signalling in striatum signals proximity and value of distant rewards. Nature 500:575–579.

Hsieh LT, Ranganath C (2015) Cortical and subcortical contributions to sequence retrieval: Schematic coding of temporal context in the neocortical recollection network. Neuroimage 121:78–90.

Hsieh LT, Gruber MJ, Jenkins LJ, Ranganath C (2014) Hippocampal activity patterns carry information about objects in temporal context. Neuron 81:1165–1178.

Kalm K, Norris D (2014) The representation of order information in auditory-verbal short-term memory. J Neurosci 34:6879–6886.

Kalm K, Norris D (2017) Reading positional codes with fMRI: Problems and solutions. PLoS ONE 12:e0176585.

Lerner Y, Honey CJ, Silbert LJ, Hasson U (2011) Topographic mapping of a hierarchy of temporal receptive windows using a narrated story. J Neurosci 31:2906–2915.

Li S, Ostwald D, Giese M, Kourtzi Z (2007) Flexible coding for categorical decisions in the human brain. J Neurosci 27:12321–12330.

Liu Y, Dolan RJ, Kurth-Nelson Z, Behrens TEJ (2019) Human Replay Spontaneously Reorganizes Experience. Cell 178:640–652 e614.

Ma L, Hyman JM, Phillips AG, Seamans JK (2014) Tracking progress toward a goal in corticostriatal ensembles. J Neurosci 34:2244–2253.

Mitchell DJ, Bell AH, Buckley MJ, Mitchell AS, Sallet J, Duncan J (2016) A Putative Multiple-Demand System in the Macaque Brain. J Neurosci 36:8574–8585.

Nili H, Wingfield C, Walther A, Su L, Marslen-Wilson W, Kriegeskorte N (2014) A toolbox for representational similarity analysis. PLoS Comput Biol 10:e1003553.

Preston AR, Eichenbaum H (2013) Interplay of hippocampus and prefrontal cortex in memory. Curr Biol 23:R764–773.

Radvansky GA, Zacks JM (2017) Event Boundaries in Memory and Cognition. Curr Opin Behav Sci 17:133–140.

Ranganath C, Ritchey M (2012) Two cortical systems for memory-guided behaviour. Nat Rev Neurosci 13:713–726.

Reagh ZM, Ranganath C (2018) What does the functional organization of cortico-hippocampal networks tell us about the functional organization of memory? Neurosci Lett 680:69–76.

Reverberi C, Gorgen K, Haynes JD (2012) Distributed representations of rule identity and rule order in human frontal cortex and striatum. J Neurosci 32:17420–17430.

Robin J, Moscovitch M (2017) Details, gist and schema: hippocampal-neocortical interactions underlying recent and remote episodic and spatial memory. Curr Opin Behav Sci 17:114–123.

Rosenbaum DA, Kenny SB, Derr MA (1983) Hierarchical control of rapid movement sequences. J Exp Psychol Hum Percept Perform 9:86–102.

Schneider DW, Logan GD (2006) Hierarchical control of cognitive processes: switching tasks in sequences. J Exp Psychol Gen 135:623–640.

Shidara M, Richmond BJ (2002) Anterior cingulate: single neuronal signals related to degree of reward expectancy. Science 296:1709–1711.

Sigala N, Kusunoki M, Nimmo-Smith I, Gaffan D, Duncan J (2008) Hierarchical coding for sequential task events in the monkey prefrontal cortex. Proc Natl Acad Sci U S A 105:11969–11974.

Smith V, Mitchell DJ, Duncan J (2018) Role of the Default Mode Network in Cognitive Transitions. Cereb Cortex 28:3685–3696.

Speer NK, Zacks JM, Reynolds JR (2007) Human brain activity time-locked to narrative event boundaries. Psychol Sci 18:449–455.

Sridharan D, Levitin DJ, Chafe CH, Berger J, Menon V (2007) Neural dynamics of event segmentation in music: converging evidence for dissociable ventral and dorsal networks. Neuron 55:521–532.

Walther A, Nili H, Ejaz N, Alink A, Kriegeskorte N, Diedrichsen J (2016) Reliability of dissimilarity measures for multi-voxel pattern analysis. Neuroimage 137:188–200.

Wen T, Mitchell DJ, Duncan J (2019) The functional convergence and heterogeneity of social, episodic, and self-referential thought in the default mode network. bioRxiv.

Woolgar A, Hampshire A, Thompson R, Duncan J (2011) Adaptive coding of task-relevant information in human frontoparietal cortex. J Neurosci 31:14592–14599.

Yeo BT, Krienen FM, Sepulcre J, Sabuncu MR, Lashkari D, Hollinshead M, Roffman JL, Smoller JW, Zollei L, Polimeni JR, Fischl B, Liu H, Buckner RL (2011) The organization of the human cerebral cortex estimated by intrinsic functional connectivity. J Neurophysiol 106:1125–1165.

Zacks JM, Tversky B (2001) Event structure in perception and conception. Psychol Bull 127:3–21.

Zacks JM, Braver TS, Sheridan MA, Donaldson DI, Snyder AZ, Ollinger JM, Buckner RL, Raichle ME (2001) Human brain activity time-locked to perceptual event boundaries. Nat Neurosci 4:651–655.

Zylberberg A, Dehaene S, Roelfsema PR, Sigman M (2011) The human Turing machine: a neural framework for mental programs. Trends Cogn Sci 15:293–300.

