## Extended Data for "Hierarchical representation of multi-step tasks in multiple-demand and default mode networks"

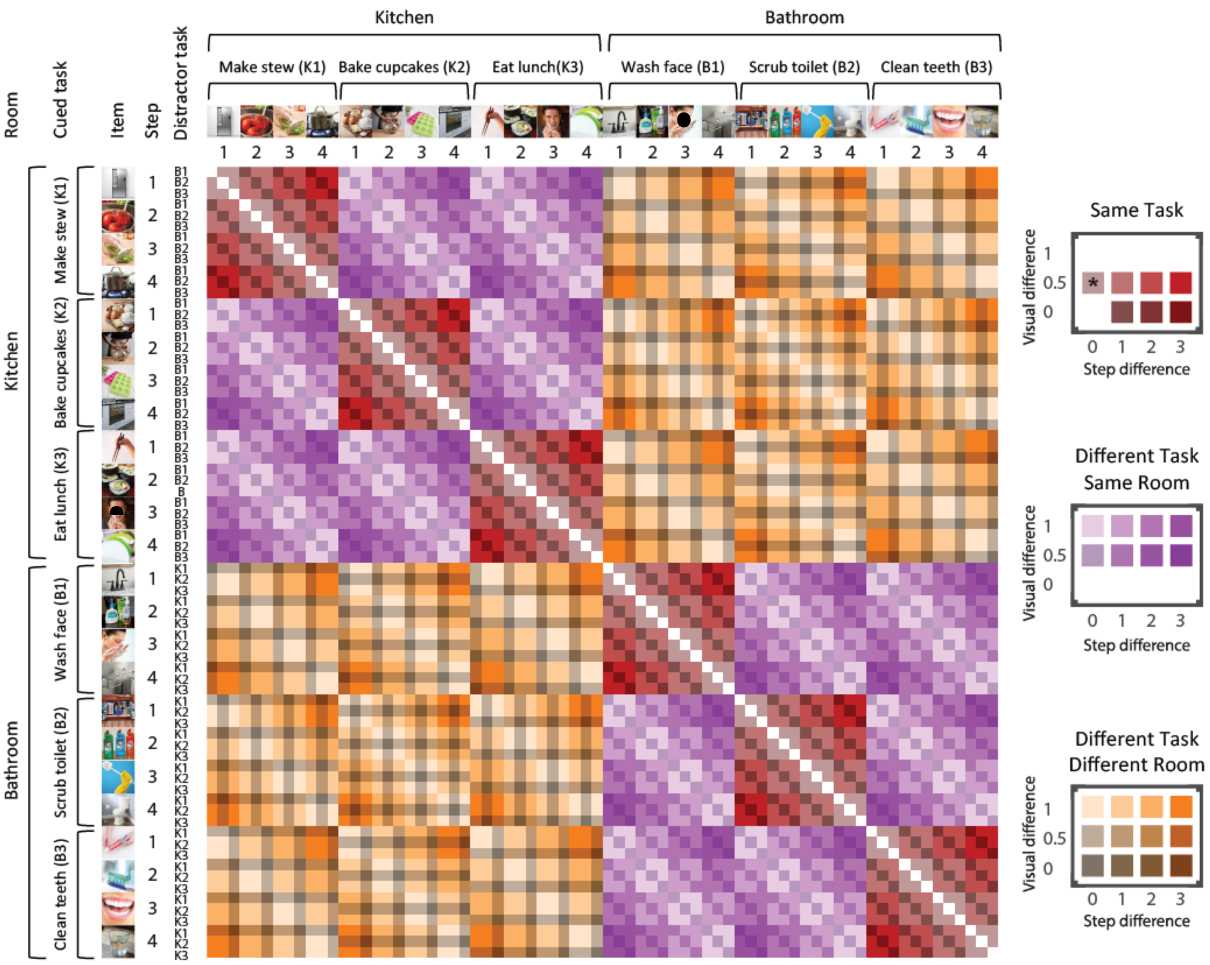


*Figure 2-1. Full representational similarity model. LDC dissimilarities were computed between every possible pair of event types (6 cued tasks × 4 steps × 3 distractor tasks per cued task), generating a 72 × 72 RDM. Diagonal cells of the RDM are zero by definition as they do not reflect a dissimilarity between different events. Off-diagonal cells reflect pattern dissimilarity between events that differ in room, cued task, step, item, and/or visual difference. These included event pairs that shared the same cued task (red cells), shared the same room but different cued task (purple cells), or differed in both room and cued task (orange cells). Saturation indicates the difference in steps between event pairs, and brightness indicates the difference in the possible stimuli presented in the visual search arrays (see main text). In the distractor task column, the six tasks are labeled K1, K2, K3, B1, B2, and B3 (with K indicating a kitchen task and B indicating a bathroom task). In the upper right panel, the asterisk indicates the only case where item is matched; this was excluded from the model fit so that the intercept could be interpreted as item coding. Images of faces are covered to comply with bioRxiv requirements.*

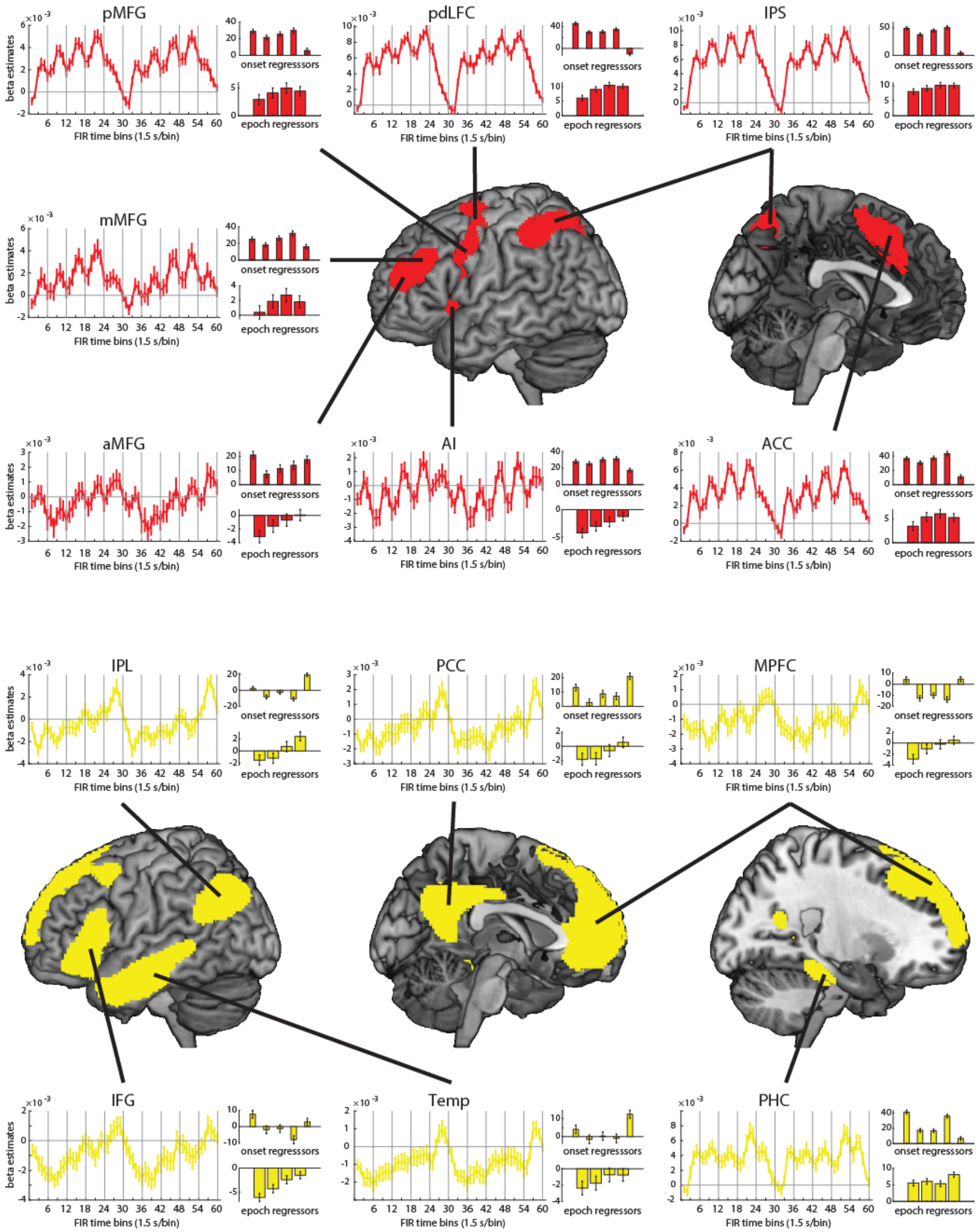


*Figure 4-1. FIR time-courses and activation profiles of onset and epoch responses in individual ROIs in the MD network (red) and DMN (yellow). The layout is the same as Figure 4. aMFG/mMFG/pMFG: anterior, middle, and posterior middle frontal gyrus; pdLFC: posterior-dorsal lateral frontal cortex; IPS: intraparietal sulcus; AI: anterior insula; ACC: anterior cingulate cortex; IPL: inferior parietal lobule; PCC: posterior cingulate cortex; MPFC: medial prefrontal cortex; IFG: inferior frontal gyrus; Temp: lateral/anterior temporal cortex; PHC: parahippocampal cortex.*

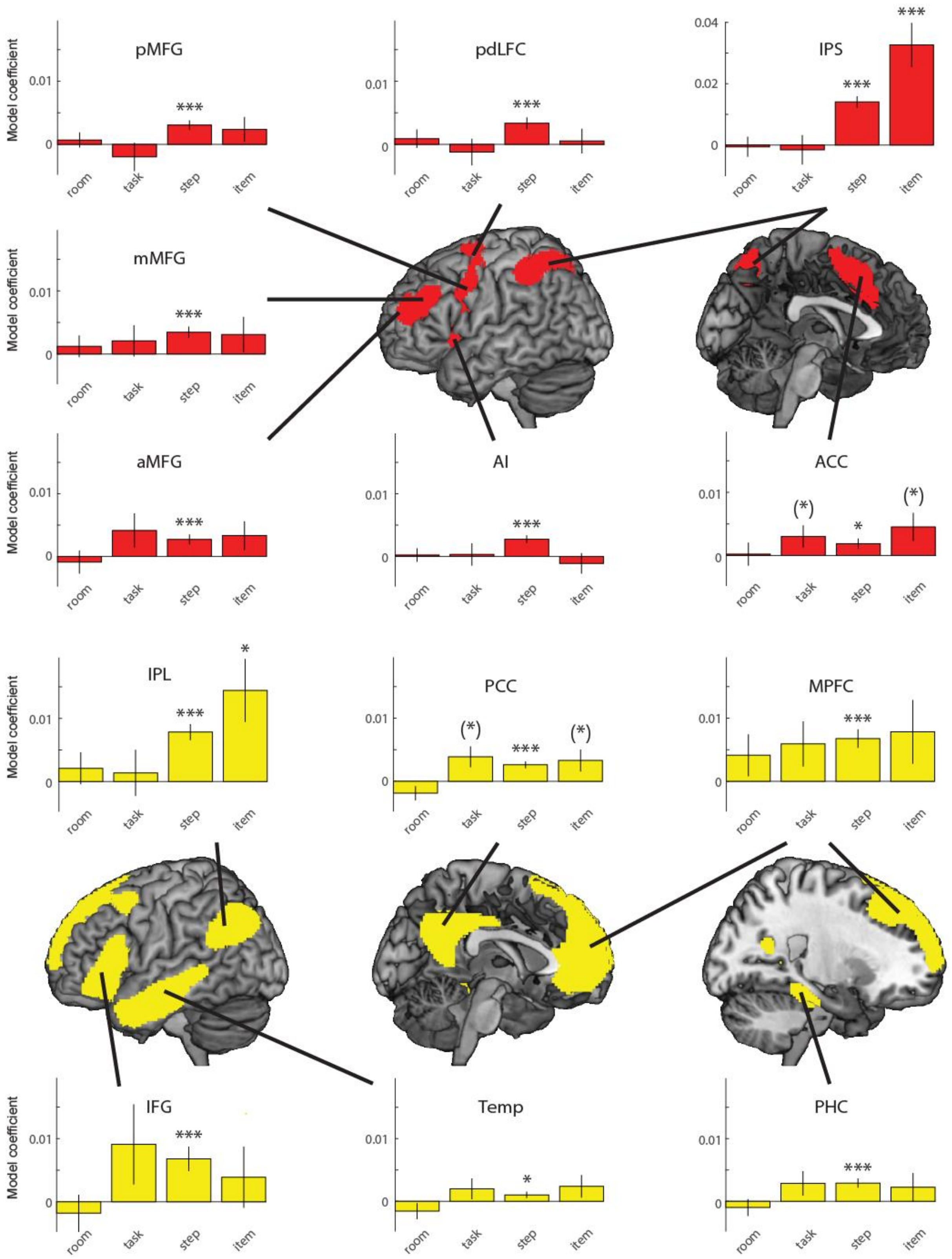


*Figure 6-1. Coding of room, task, step, and item information in individual ROIs in the MD network (red) and DMN (yellow). Asterisks indicate significance of 1-tailed t-tests against zero, after FDR-correction across the number of ROIs, separately for each information type; *** p < 0.001, ** p < 0.01, * p < 0.05. Additionally, (*) indicates significance of p < 0.05 before FDR correction. Error bars represent +/- 1 standard error of the mean across subjects. ROIs are the same as in Figure 4-1.*

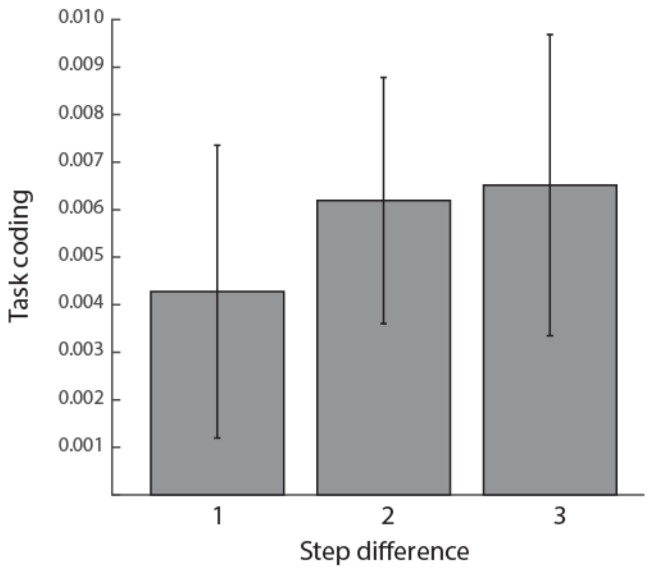


*Figure 6-2. Task coding as a function of step difference in the DMN network ROI. Error bars represent +/- 1 standard error of the mean across subjects.*
